# Impact of oral *Chlamydia* Vaccination on Host Gut Microbiome and Metabolites Composition

**DOI:** 10.1101/2024.11.11.623094

**Authors:** Youyou Huang, Jiao Wan, Chuqiang Shu, Qi Zhang, Zengzi Zhou, Xin Sun, Luying Wang, Tianyuan Zhang, Qi Tian

## Abstract

*Chlamydia trachomatis*, an intracellular pathogen, is recognized as the most common sexually transmitted bacterial infection among women worldwide. *Chlamydia* infections can lead to undesirable clinical outcomes, including pelvic inflammatory disease and infertility. Recently, the gut has been identified as a niche for Chlamydia colonization; however, despite the biological impact on the host remaining under investigation, oral inoculation of Chlamydia as a whole-organism vaccine has been reported as a promising strategy for preventing genital Chlamydia infections. Few studies have evaluated the impact of oral Chlamydia vaccination on the gut microbiome and metabolite changes. In this study, we assessed time-series alterations in the gut microbiome and metabolites following oral *Chlamydia* inoculation, and we analyzed the composition and correlation between serum immune parameters and the sequencing profiles in the host. We identified 129 microbial changes and 186 significantly different metabolites in the gut across various vaccination approaches during the 30-day immunization process. Additionally, we discussed potential biomarkers of effective immunization based on correlation analysis.

**IMPORTANCE:** *Chlamydia* infections primarily lead to morbidity rather than mortality. Consequently, in developing and implementing a *Chlamydia* vaccine, the utmost priority is ensuring safety. As a promising yet controversial approach, live oral vaccination for *Chlamydia* raises concerns regarding its impact on the host’s gut environment. Our study not only investigates changes in the gut microbiome and metabolites during vaccination but also identifies potential biomarkers during immunization. These findings are crucial for the development of whole-organism oral *Chlamydia* vaccines and provide valuable insights into the long-term colonization of *Chlamydia* in the gut.

## INTRODUCTION

*Chlamydia trachomatis* (*C. trachomatis*) is the most common sexually transmitted infection globally, and it can lead to serious complications in the female upper genital tract, such as pelvic inflammatory disease and infertility(1). Although antibiotics can effectively treat the infection, the asymptomatic nature of infection poses challenges for prevention and timely intervention(2). In this context, vaccination could play a crucial role in preventing *C. trachomatis* infections and offering a long-term solution to this challenge(3). Currently, there are few licensed *Chlamydia* vaccines available for human use(4). Among the various types of vaccines, whole-organism *Chlamydia* vaccines have shown effective protection in animal studies(5). In 2015, a deactivated form of *C. trachomatis* was modified and used to induce protective immunity in preclinical studies(6). Additionally, live-attenuated vaccines for *C. psittaci* and *C. abortus* have been approved for the protection of animals against Chlamydia infections(7, 8).

Studies have demonstrated that oral inoculation of live *Chlamydia* induces robust transmucosal protection in mice, resulting in a shortened course of *Chlamydia* reinfection and reduced pathology in the upper genital tract(9). Despite the efficacy of *Chlamydia* vaccination in the gut, the safety of the vaccine still requires thorough investigation. While research suggests that gut inoculation with live *Chlamydia* does not cause visible pathology in either the gut or the genital tract, there remains a concerning possibility that live organisms could recover from the immunized host.

As a genital pathogen, *Chlamydia* is often detected in the gastrointestinal (GI) tracts of various hosts, including humans(10, 11). Studies indicate that the gut may serve as a natural niche for long-term *Chlamydia* colonization, and some findings suggest that gut *Chlamydia* co-infection with genital *Chlamydia* during the acute phase could potentially trigger pathogenic immune responses in the upper genital tract(12, 13). Research has examined the potential of using inactivated *Chlamydia* as a vaccine to induce protective immunity while reducing the risks associated with live *Chlamydia*.

However, findings suggest that inactivated *Chlamydia* may offer inferior protective immunity relative to live *Chlamydia*. This discrepancy may arise from the ability of live *Chlamydia* to replicate and disseminate, which enhances the diversity of antigens presented to the immune system. The reasons behind the superior protective immunity induced by live *Chlamydia* compared to killed organisms remain unclear(4). Investigating the metabolic changes and patterns that occur following the inoculation of live versus deactivated *Chlamydia* could provide valuable insights.

Recent research indicates that the need for a viable *Chlamydia* vaccine to generate protective immunity may be circumvented by employing adjuvant-formulated nanoparticles, which have been associated with the enhancement of transmucosal tissue-resident memory T cell responses(6). Notably, tissue-resident memory T cell responses are significantly influenced by the metabolites generated from host-pathogen interactions. This underscores the necessity of investigating the gut microbiome and the profiles of metabolites during the immunization process. Understanding these relationships can provide insights into how gut microbes and their metabolic byproducts contribute to the efficacy of immune responses, potentially guiding the development of more effective vaccine strategies.

In this study, we assessed the changes in microbial populations and metabolite profiles over time following oral inoculation with live or deactivated *Chlamydia* EB. Additionally, we evaluated the resulting immune responses and explored the correlations between microbial dynamics and metabolite levels post-immunization.

## RESULTS

### Oral inoculation with live *Chlamydia* results in sustained gut colonization and induces protective immune response

Three groups of mice were intragastrically inoculated with live *Chlamydia*, deactivated *Chlamydia*, and SPG as a vehicle control, respectively. The infection burden was monitored by calculating *Chlamydia* inclusions recovered from genital and rectal swabs. After inoculation, live *Chlamydia* oral inoculation resulted in persistent *Chlamydial* colonization in the gut, and live *Chlamydia* is detectable in rectal swabs more than 50 days post-inoculation. Notably, no *Chlamydia* was recovered from genital swabs, indicating no cross-contamination from the gut to the genital tract after vaccination (Fig. 1A panel e, f). The persistence of *Chlamydia* in the gut after immunization can also be monitored using in vivo imaging with Luciferin-labeled *Chlamydia* (Fig. 1C).

**Figure 1:**
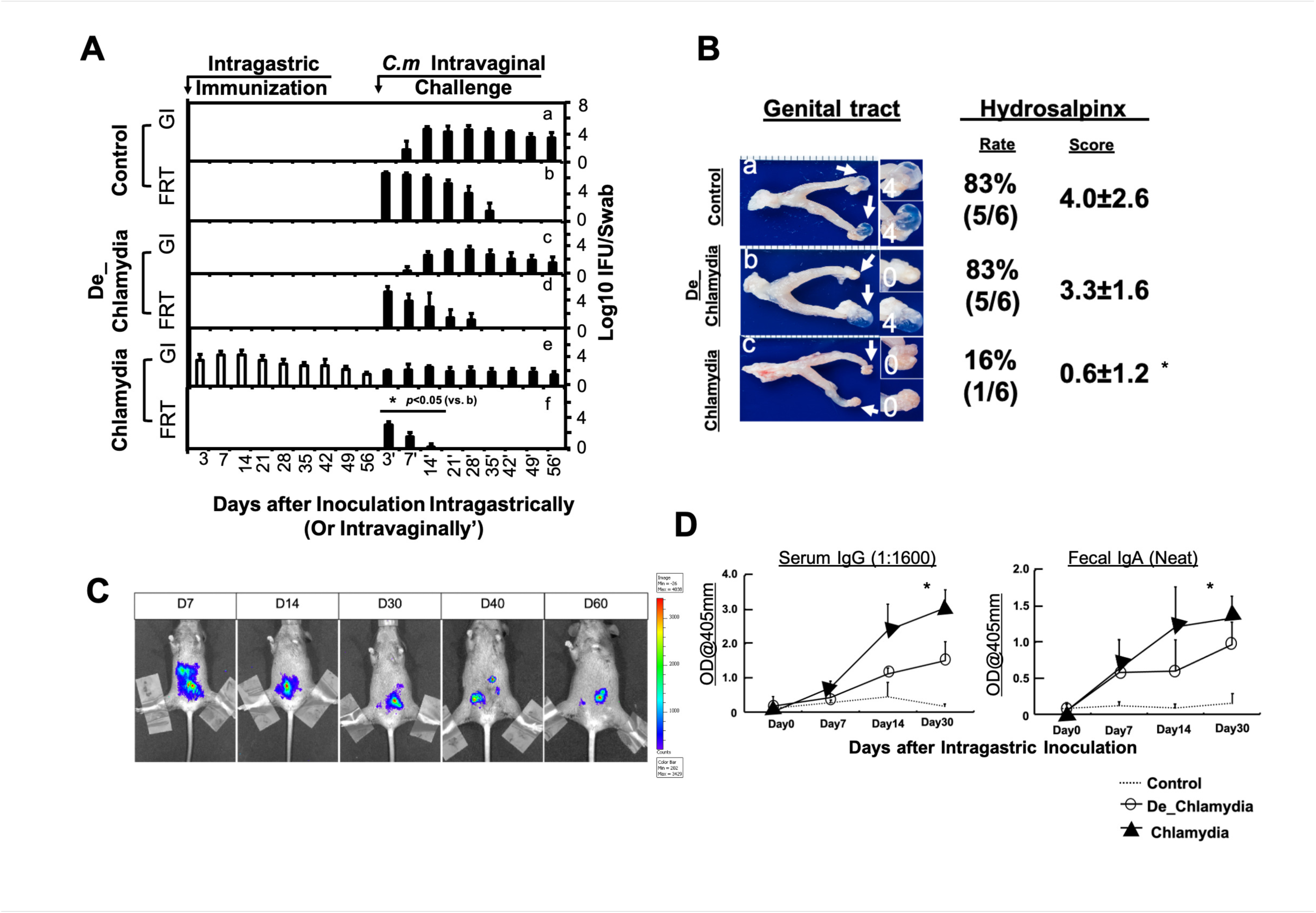
Oral administration of live *Chlamydia* leads to a long-lasting gut infection and exerts protective immune effects against genital pathology. [A]This panel illustrates the shedding of Chlamydia from the gastrointestinal (GI) and female reproductive tracts (FRT) after vaccination in the GI tract, along with an assessment of infection severity among different treatment groups following inoculation and subsequent challenge. C57BL/6J mice were intragastrically inoculated with SPG buffer only (control group, n=6) (a and b), 2 × 10^5 IFU of deactivated *C. muridarum* (immunization group 1, n=6) (b and c), or 2 × 10^5 IFU of live *C. muridarum* (immunization group 2, n=6) (c and d). These mice were challenged intravaginally on day 56 with 2 × 10^5 IFU of *C. muridarum*. Results are presented as the log10 number of IFU per swab specimen on the y-axis. Notably, on days 3, 7, and 14 following the intravaginal challenge after oral live *Chlamydia* immunization, the immunized mice exhibited a 1,000-fold reduction in the number of IFUs, as determined by vaginal swab analysis at each time point (*p* <0.05, Wilcoxon rank-sum test). Furthermore, the overall shedding duration was significantly lessened (\**p*<0.05, Wilcoxon rank-sum test, AUC for panel f vs. panel b). [B]Mice were sacrificed on day 56 following the intravaginal challenge to evaluate upper genital tract pathology. Representative macroscopic images of an entire genital tract from both the control (a) and immunization (b, c) groups are presented. White arrows indicate oviducts exhibiting hydrosalpinges. Enlarged images of the oviduct and ovary regions are provided on the right side of the overall genital tract images, with numbers representing hydrosalpinx scores. Both incidence and severity of hydrosalpinx were quantified using a previously published scoring method. Mice immunized with live *Chlamydia* in the GI tract showed a significantly lower incidence (*p* <0.05, Fisher’s exact test) and reduced scores (\**p* <0.05, Wilcoxon rank-sum test) when compared to control mice. [C]In vivo imaging was utilized to monitor the presence of live *Chlamydia* in the mice, revealing evidence of long-term colonization. Mice were intragastrically infected with a *C. muridarum* strain expressing a luciferase gene. Bioluminescence signals produced by luciferase were detected using whole-body imaging technology on various days post-infection, with signals illustrated in red, green, and blue colors to indicate decreasing intensity. Notably, these bioluminescence signals remained detectable until day 60. [D]Mice from three groups (live *Chlamydia*, deactivated *Chlamydia*, and controls) were assessed for fecal IgA and serum IgG levels using enzyme-linked immunosorbent assay (ELISA). The anti-*C. muridarum* IgG responses at a dilution of 1:1,600 are shown for immunized mice along with the neat solution for fecal IgA (\**p* <0.05, Wilcoxon rank-sum test) compared to the levels of other antibody types.

Sixty days after immunization, the mice were intravaginally challenged with 2×10^5 infectious units (IFU) of wild-type *C. muridarum* in all three groups. The genital infection burden was significantly lower in the live *Chlamydia* group compared to the deactivated *Chlamydia* group (Fig. 1A, panel f vs d), demonstrating a more effective protective effect against genital mucosal *Chlamydia* infection. The duration of genital infection was shorter in both the *Chlamydia* group and the deactivated *Chlamydia* group compared to the control group, suggesting that the vaccination provided mucosal protection (Fig. 1A, panels f and d vs. b).

Additionally, genital pathologies assessed 56 days post genital challenge showed a significant reduction in hydrosalpinx in the live *Chlamydia* group compared to other two groups (Fig. 1B, panels a and b vs. c, *p*<0.05). No significant changes were observed between the deactivated *Chlamydia* and Control groups. These data suggest that while deactivated *Chlamydia* immunization offers a certain, though not significant, level of protection against mucosal infection, it is less effective in preventing genital pathology outcomes. Similarly, serum IgG and fecal IgA antibodies after immunizations were significantly higher in the live *Chlamydia* group compared to both the deactivated *Chlamydia* and control groups (Fig. 1D).

### Gut *Chlamydia* long term colonization changes the ultra-microscopic structure of the colon epithelium

On day 30, three mice from each treatment group were sacrificed to assess the effects of immunization on colon epithelial cells. Transmission electron microscopy revealed that the control group displayed a dense array of microvilli in the intestinal epithelial cells (Fig. 2 a1, a2). The mitochondria were uniformly distributed, and the structures contained well-defined double membranes. Additionally, visible lamellar cristae were arranged in approximately parallel layers. In contrast, the live *Chlamydia* (Fig. 2 c1, c2) group demonstrated a significant reduction in the number of microvilli, accompanied by a marked shortening of their length, as indicated by the black arrows. Moreover, numerous swollen mitochondria were observed in the cytoplasm, characterized by reduced matrix electron density, as highlighted by the red arrows. The mitochondrial cristae were also notably increased, as shown by the yellow arrows, along with many autophagosomes indicated by the white arrows.

**Figure 2:**
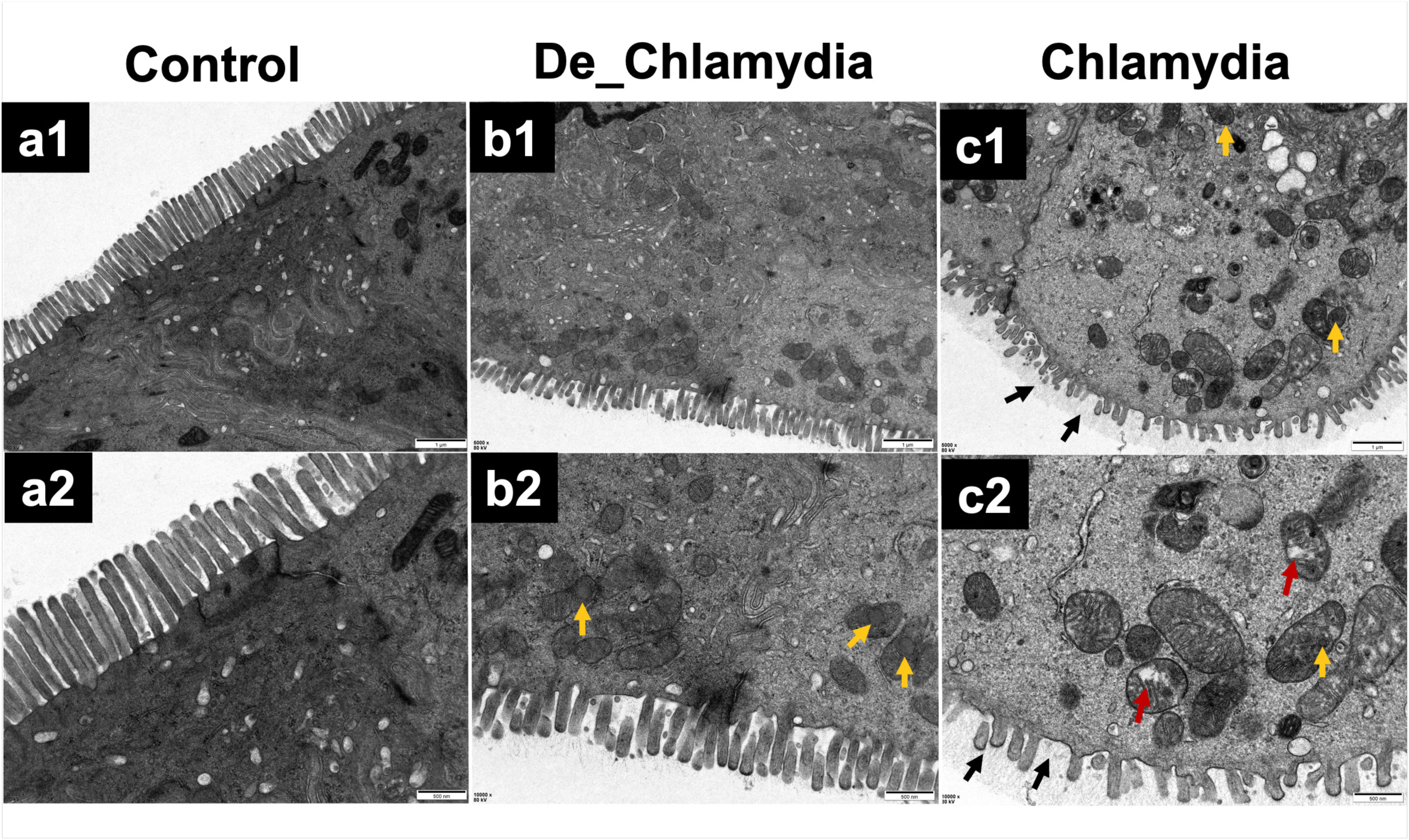
Ultrastructural analysis of colon epithelial cells following immunization. Transmission electron microscopy (TEM) images reveal variations in microvilli integrity and mitochondrial morphology of colon tissue among control, live, and deactivated *Chlamydia* treatment groups. The TEM images (×5000 and ×1000 magnifications) are presented to compare the ultrastructural characteristics of the different groups. Panels a1 and a2 depict the control group, while panels b1 and b2 represent the deactivated *Chlamydia* group, and panels c1 and c2 illustrate the live *Chlamydia* group. Yellow arrows indicate increased mitochondrial cristae, whereas red arrows highlight swollen mitochondria with reduced matrix electron density.

The group treated with deactivated *Chlamydia* (Fig. 2 b1, b2) displayed relatively shorter microvilli and an increased number of mitochondrial cristae. Despite this, significant improvements were noted when compared to the live *Chlamydia* group. Specifically, the number of microvilli in intestinal epithelial cells increased, and no swelling of mitochondria or presence of autophagosomes was observed. Additionally, immunohistochemical analyses indicated that the actively infected group exhibited a higher infiltration of inflammatory immune cells.

### Time series analysis of bacterial community changes between different vaccinations

Sixty-three fecal samples contained 20049 effective sequences from 4 time points of three treatment groups were used for OUT clustering. The three groups were: Control (SPG inoculation), De_*Chlamydia* (deactivated *Chlamydia* EB oral inoculation), and *Chlamydia* (live Chlamydia EB inoculation). Fecal samples were taken at four time points after immunization: day 0, day 7, day 14, and day 30. DNA was extracted from the fecal samples, and libraries were constructed and sequenced.

The dilution curve illustrates the sequencing depth (Fig. 3 A, B, and C), with adequate sequencing depth indicated by the plateau reached in the number of reads on the x-axis. Panel A displays the dilution curves for each sample, while Panel B presents the sample curves grouped by treatment and time point. Panel C shows the dilution curves for samples, also grouped by treatment and time point, along with their 95% confidence intervals. These findings suggest that the majority of intestinal microbial diversity was captured in this study and that the sequencing effort was sufficient to accurately represent the gut microbiota diversity of the samples.

**Figure 3:**
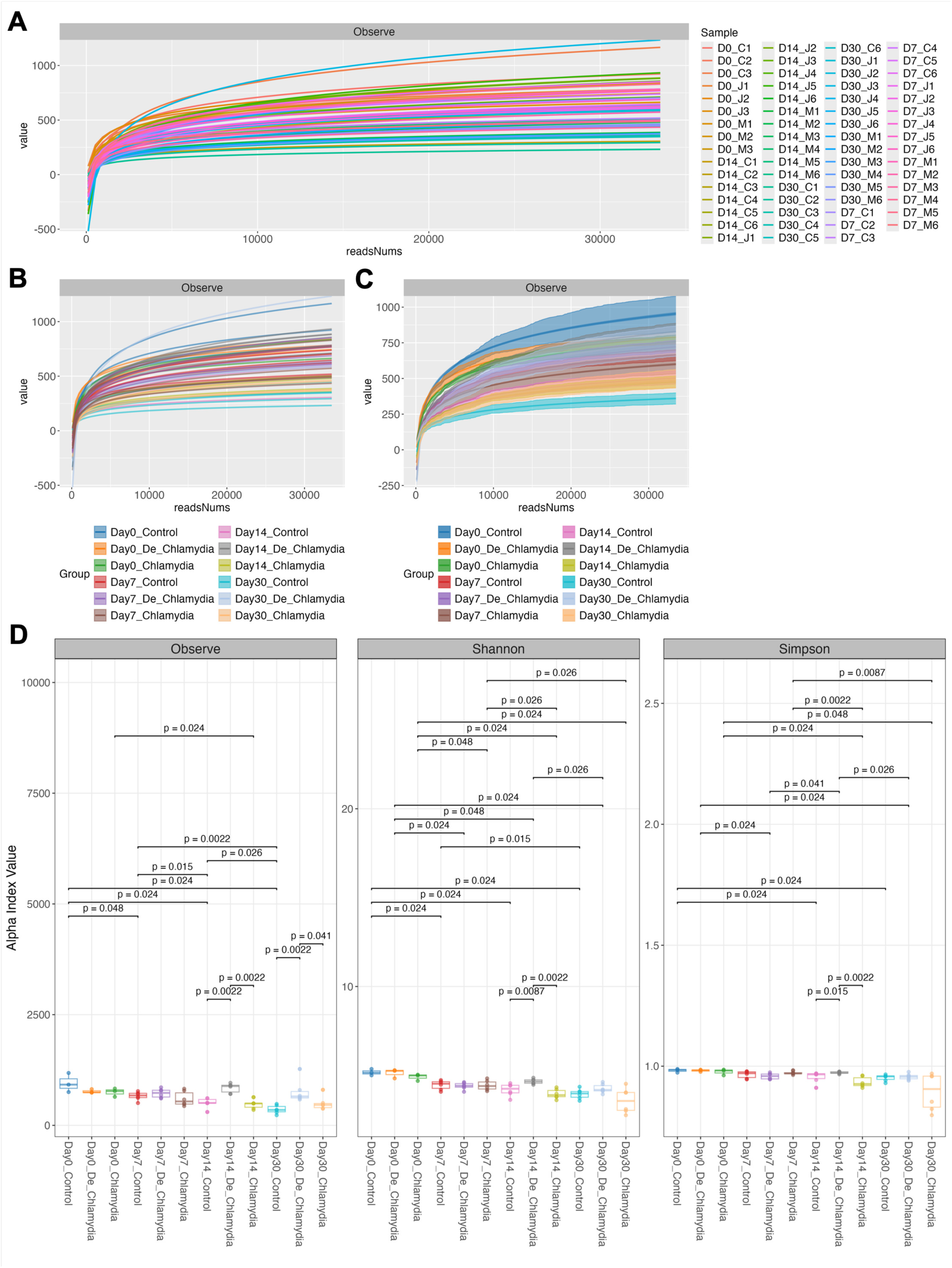
Microbial alpha diversity analysis illustrating the dynamic changes in microbial communities over the immunization time series. Panels A, B, and C present dilution curves for various samples and treatment groups, confirming that adequate sequencing depth was achieved. The dilution curves reflect sequencing depth information, with the x-axis representing the number of reads. It is generally accepted that the sequencing depth is considered adequate when the x-axis reads reach a plateau. Specifically, panel A shows the dilution curves for each sample; panel B displays the sample curves grouped by treatment and time points; and panel C presents the dilution curves (with 95% confidence intervals) grouped by treatment and time points. Panel D summarizes the α-diversity indices across groups at different time points, evaluating Observed species, Shannon index, Simpson index (D), Chao1, ACE, and Pielou’s evenness index (Fig. S1). Only p-values less than 0.05 are indicated, as determined by the Wilcoxon rank-sum test.

The α-diversity indices, including Observed Species, Shannon, and Simpson indices (Fig.3 D) and others (Fig. S1), were utilized to evaluate the diversity and richness of the GI flora after inoculations. These indices significantly decreased in both the Control and *Chlamydia* groups over time, whereas the De_*Chlamydia* group exhibited an increase on day 14. At each time point, the indices were similar between the Control and Chlamydia groups, while the De_*Chlamydia* group demonstrated higher indices on days 14 and 30.

The figure 4 illustrates the dynamic changes in phyla and genus -level flora abundance over time, both in grouped and individual samples. The dominant phyla included *Bacteroidetes*, *Firmicutes*, *Proteobacteria*, and *Verrucomicrobia*, with fluctuations in their abundance observed over time and differing between groups (Fig. 4 A, B). And the dominant genus identified included *S24-7*, *Prevotella*, *Lactobacillus*, *Akkermansia*, *Lachnospiraceae*, *Clostridiales* along with other genera (Fig. 4 C, D). The genus level of *Firmicutes*/*Bacteroidetes* (Fir/Bac) ratios were analyzed for each group, revealing that the *Chlamydia* group maintained a relatively stable Fir/Bac ratio, while the other groups exhibited an increased Fir/Bac ratio by day 30 (Fig. 4 E, F).

**Figure 4:**
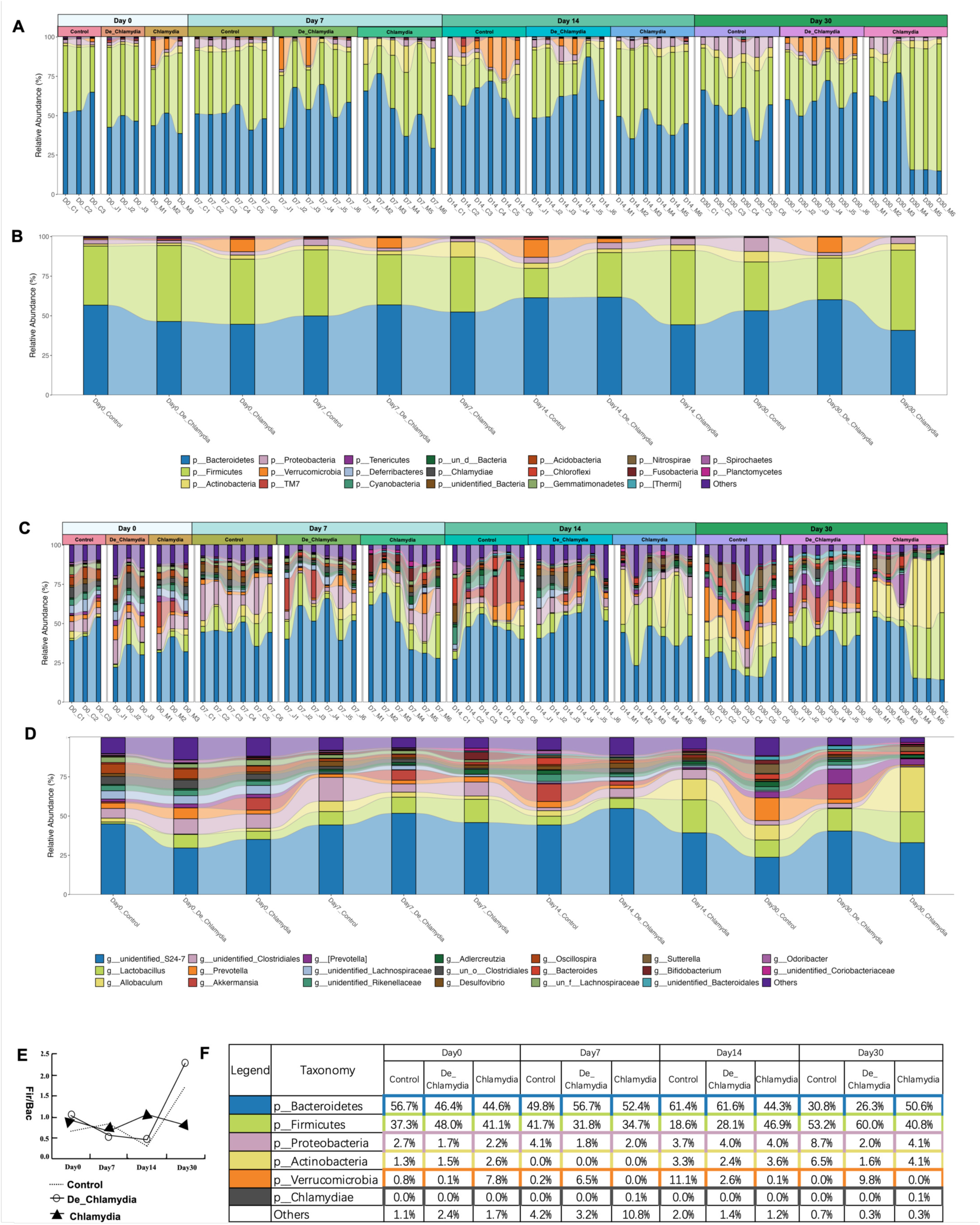
Temporal changes in gut microbiota composition and abundance at the phyla and genus levels. Panels A and B illustrate the top 20 dominant phyla and their fluctuations across samples and treatment groups over the time series. Panels C and D highlight the top 20 changes in specific genera among the samples and combined groups. Panel E presents the Firmicutes to Bacteroidetes (Fir/Bac) ratio among the groups throughout the time series. Finally, Panel F features a table detailing the top five dominant phyla and the detection of *Chlamydia* during the time series across the different groups by 16S rRNA sequencing.

To further investigate the impact of oral *Chlamydia* vaccination on the gut flora community, β-diversity analysis was conducted. Principal Coordinates Analysis (PCoA) of weighted and unweighted UniFrac distance matrices demonstrated distinct clustering of bacterial microbiota between different 12 groups (adnois R^2^=0.49, *p*=1e-04, anosim R= =0.0.66, *p*=1e-04) (Fig. 5, A, B). Bray-Curtis dissimilarity analysis was conducted to assess the differences in β-diversity between various time points and groups, as shown in Figure 5 (C). On days 14 and 30, the β-diversity of the De_*Chlamydia* group was significantly different from that of the Control and *Chlamydia* groups.

**Figure 5:**
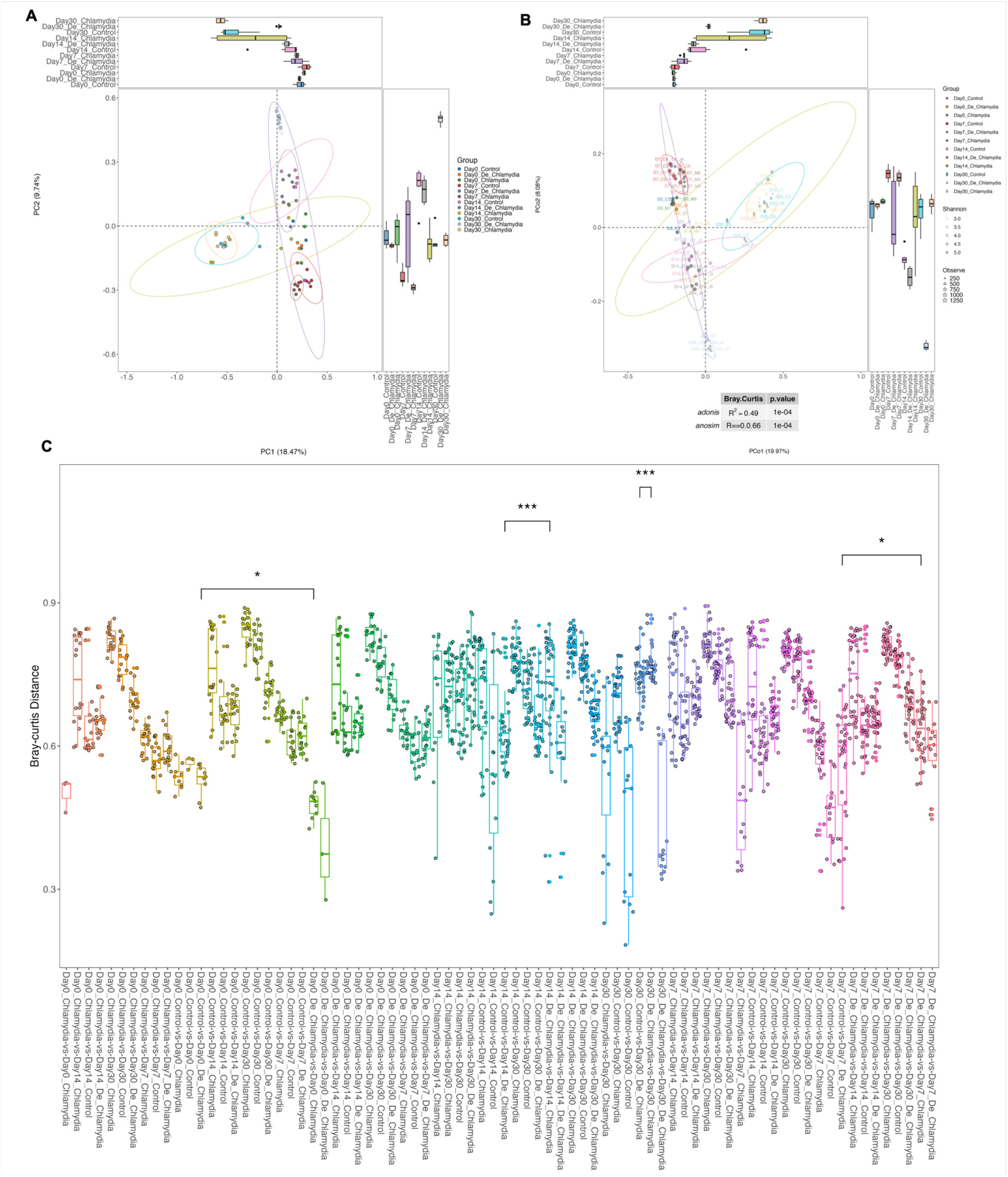
β-diversity analysis of gut microbial communities among treatment groups over time. Panel A illustrates Principal Component Analysis (PCA), an unsupervised dimensionality reduction method and Panel B presents Principal Coordinates Analysis (PCoA), a supervised dimensionality reduction method that identifies distinct clustering of microbial communities across groups. The Bray-Curtis dissimilarity is utilized to evaluate the compositional differences in species across groups (Panel C), based on quantitative characteristics of the species present in the samples (\**p* < 0.05, \*\*\**p* < 0.001)

Additionally, we identified major alterations in gastrointestinal bacterial taxa through sample and group LEfse analysis (Fig. 6, A, B), and detected 129 significantly changed GI taxa abundances over the time series (LDA Score > 2.0, Fig. S4). Notably, at the genus level, *Bifidobacterium* and *Chlamydia* were enriched in the *Chlamydia* group on day 7. The genera *Allobaculum*, *Lactobacillus*, and g_*Clostridium_*f_*Clostridiaceae* showed increased abundance in the *Chlamydia* group on days 14 and/or 30. In the De_*Chlamydia* group, a rise in the abundance of genera *Yaniella*, *Corynebacterium*, and *Staphylococcus* was also observed on day 14. Across all three groups,the g_*Clostridium_*f_*Lachnospiraceae* exhibited a decreasing trend over the time series, while *Lactobacillus* demonstrated an increasing trend(Fig. 6, C).

**Figure 6:**
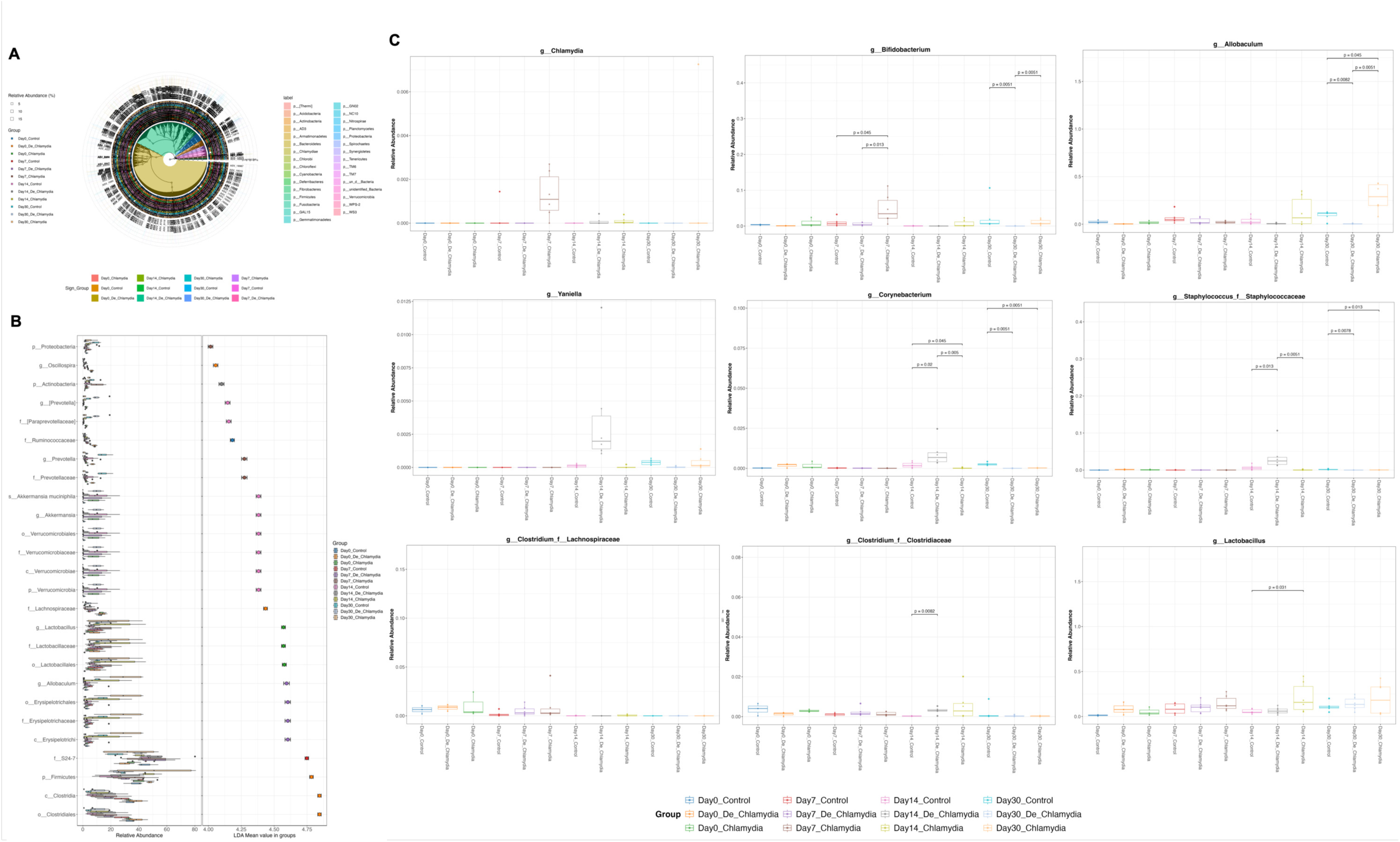
LEfSe analysis identifying differentially abundant taxa in the gut microbiome during vaccination. Panel A visualizes the group LEfSe results through a microbial phylogenetic graph (Kruskal-Wallis test, *p*-values adjusted with the False Discovery Rate (FDR), and post hoc analysis performed with the Wilcoxon test, applying a cutoff of *p* < 0.05). Panel B displays microbes with Linear Discriminant Analysis (LDA) scores greater than 4 (for those with LDA > 2, see Fig. S4). Panel C highlights representative microbes identified and significant changes in microbial taxa abundance at the genus level over time are noted (\**p <* 0.05, Wilcoxon test).

### Metabolic alterations analysis over the time series revealed unique changes in metabolites in the live *Chlamydia* group on day 7

The metabolites in the time series were obtained from LC-MS analysis (Fig. 7). We analyzed the intersecting differences in metabolites among different groups over the time series, resulting in the preliminary identification of 1,029 differential metabolites in negative mode and 1,362 in positive mode. To account for the dynamic changes of metabolites over time, we established a time series for the expression of the identified characteristic peaks and performed Mfuzz analysis to identify clusters with opposite expression trends. Ultimately, we observed significant differences between cluster 4 and cluster 6 in the negative mode and between cluster 1 and cluster 6 in the positive mode, all of which were from the *Chlamydia* group on Day 7 (Fig. 7, A). We extracted the data from these clusters for further analysis. By intersecting clusters from the Mfuzz analysis with the previously identified metabolites, we obtained a set of 295 metabolites (Fig. 7, B) from negative mode and 395 metabolites from positive mode that exhibited notable changes in expression over the time series.

**Figure 7:**
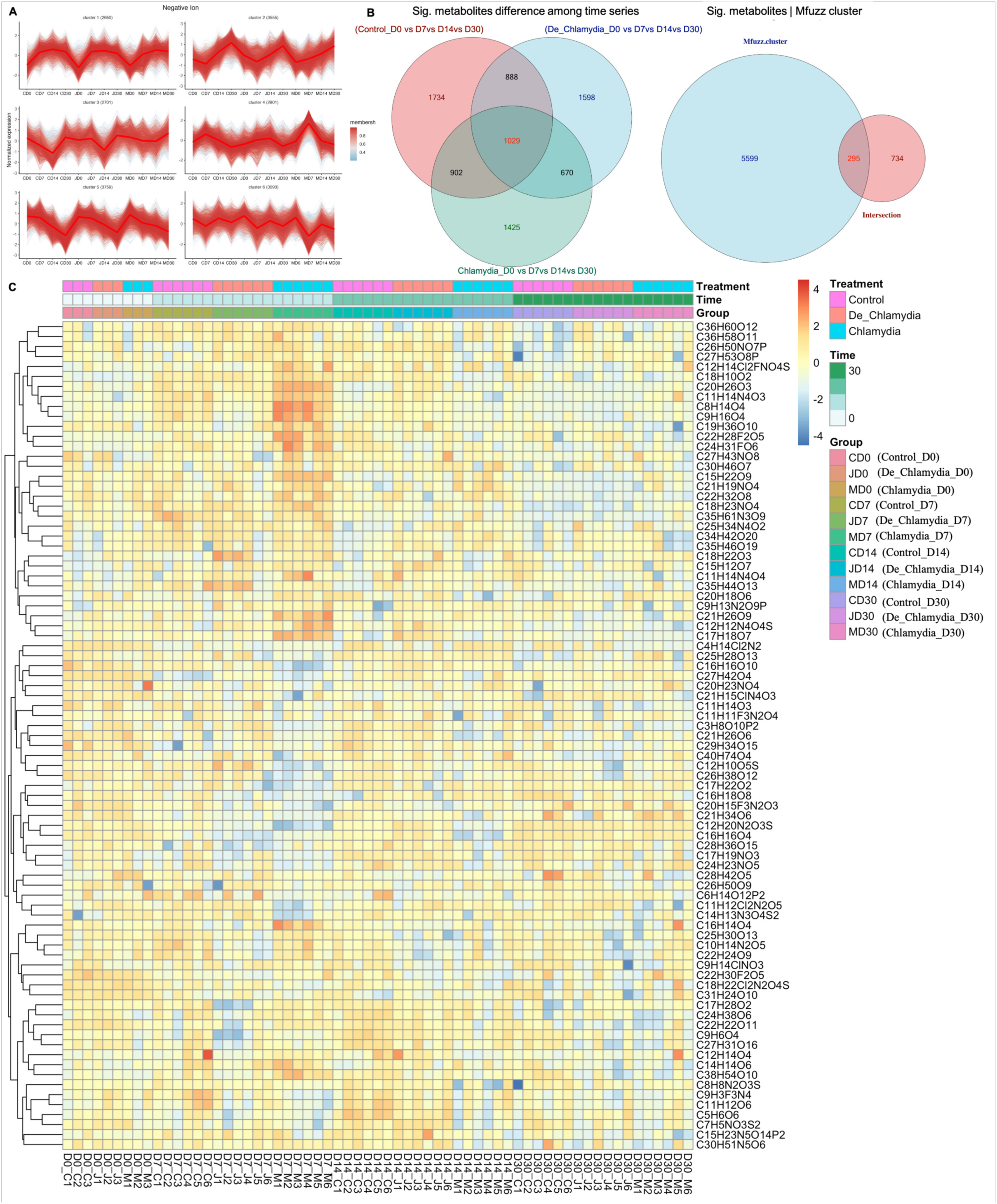
LC-MS analysis of metabolite changes in response to oral *Chlamydia* vaccination. Differential Metabolites identified in negative ion modes are outlined, with clusters illustrating expression trends over time. Fecal samples were collected at different time points (D0, D7, D14, D30) alongside feces for 16S rRNA sequencing and the fecal samples were subjected to LC-MS analysis. [A] The metabolomic profiles from various time points and groups were analyzed using the Mfuzz method. This analysis aimed to identify clusters of metabolite patterns across different samples and to establish relationships between time and metabolites within these clusters. Notably, on Day 7 (Clusters 4 and 6), the live Chlamydia group exhibited the most significant metabolic changes among all groups and time points. [B]We focused on the differential metabolites produced at each time point after each treatment, then considered the intersections of three different treatments to identify biomarkers that may vary with time across each condition. The analysis of differential metabolites included comparisons: CD0 vs. CD7 vs. CD14 vs. CD30, JD0 vs. JD7 vs. JD14 vs. JD30, and MD0 vs. MD7 vs. MD14 vs. MD30 (Supplemental table2). An intersection of the differential substances yielded 1,029 metabolite entries. [C] By intersecting the metabolites from Mfuzz clusters (Clusters 4 and 6) with these 1,029 substances, we identified 295 metabolites that exhibited significant changes over time and showed differences across groups with *p < 0.05*. As depicted in the figure, the final intersection comprised 295 metabolites, which displayed notable fluctuations at certain time points and demonstrated significant differences in comparisons across all groups. A matching analysis of these 295 metabolites with the company-provided metabolite identification results and resulted in the identification of 83 specific compounds

Subsequently, we matched these metabolites with the first and second-level identification results provided by the company (Supplemental table1), successfully identifying 83 specific metabolites from negative mode and 103 specific metabolites from positive mode (Fig. S5).

### Potential associations among the GI microbiome, metabolites, and *Chlamydia* specific antibody

We conducted a Pearson correlation analysis to explore the relationships between 129 predominant bacterial species from gut microbiome analysis (LDA > 2), 186 metabolites identified through LC-MS analysis (negative mode 83 and positive mode 103), and levels of *Chlamydia*-specific antibodies (Serum IgG and Fecal IgA) at various time points (Fig. S6). This analysis enabled us to identify both positively and negatively correlated bacterial communities, metabolites, and antibodies with a significance level of P < 0.05. Based on the biological significance and existing scientific literature concerning infection and immunity, we illustrated these relationships using heat maps, highlighting the most significant correlations for presentation and future discussion (Fig. 8).

**Figure 8:**
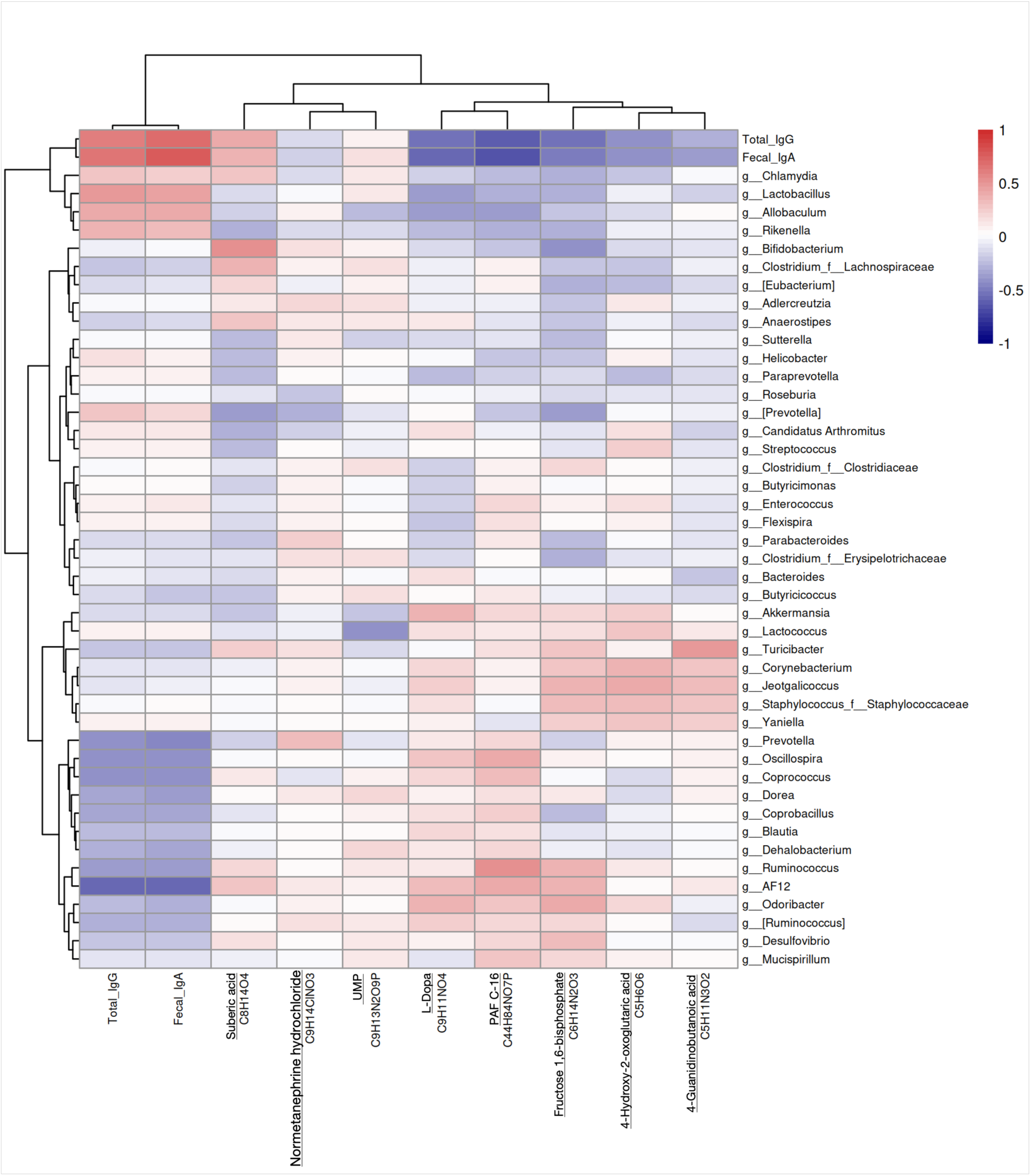
Pearson correlation analysis of the gut microbiome, metabolites, and *Chlamydia*-specific antibodies. The heat map illustrates significant relationships among microbial taxa (at the genus level), metabolite levels, and indicators of immune response. A total of 129 dominant microbial taxa (LDA > 2) were analyzed in correlation with 186 metabolites identified from the non-targeted metabolomics approach (comprising 83 negative and 103 positive metabolites), as well as serum IgG and fecal IgA levels from different time points. The figure displays microbes at the genus level and metabolites that exhibit biological relevance.

Key findings reveal a negative correlation between the production of protective antibodies (Total IgG and Fecal IgA) and the genus *AF12* (from the *Rikenellaceae* family), with Pearson values of -0.56 and -0.55, respectively. In addition to *AF12*, genera such as *Prevotella*, *Oscillospira*, *Coprococcus*, *Blautia*, *Dehalobacterium*, *Ruminococcus*, *Odoribacter*, and *Desulfovibrio* also displayed negative correlations with antibody levels. Furthermore, these antibodies were negatively associated with metabolites like L-Dopa and PAF C-16, with Pearson values ranging from -0.51 to -0.63. Conversely, genera including *Chlamydia*, *Lactobacillus*, *Allobaculum*, and *Rikenella* showed positive correlations with antibody production, exhibiting Pearson values between 0.25 and 0.48.

## DISCUSSION

Chlamydia is a sexually transmitted infection that can lead to severe female genital sequelae(1). Studies have shown that Chlamydia can ascend to the upper female genital tract and infect tubal epithelial cells, potentially resulting in tubal damage, adhesion, fibrosis, or hydrosalpinx (source: [CDC](http://www.cdc.gov/std/tg2015/chlamydia.htm)). The pathogenic mechanisms of Chlamydia infection are still under investigation, and the persistence and elusive nature of the pathogen pose challenges for infection control.(2) In recent years, the gastrointestinal (GI) tract has begun to be recognized as a niche for Chlamydia colonization, yet its biological effects remain largely unknown. Animal studies have indicated that Chlamydia can persistently colonize the GI tract, and pre-inoculation of Chlamydia in the GI tract of naïve mice may induce a transmucosal protective immune response to future Chlamydia infections(9, 14). Similarly, Chlamydia is routinely detected in the human GI tract according to numerous clinical studies(10). C. trachomatis can spread to the GI tract through sexually dependent pathways, regardless of oral or anal sex history(15). Research indicates that the GI tract can serve as a final reservoir for persistent Chlamydia infections, regardless of the various routes of infection into the host.

Chlamydia is believed to inhabit the gut through mechanisms that involve the evasion of host immune responses and the exploitation of plasmid-related resistance against the killing mechanisms of the GI tract(16). Research suggests that the process of Chlamydia spreading to and colonizing the GI tract occurs by steps(17). Certain mutants of *C. muridarum* display reduced capabilities in colonizing the GI tract(18, 19); for instance, mutants lacking plasmid functions exhibit significant difficulties in colonizing the upper GI tract, whereas those deficient in specific chromosomal genes show more pronounced obstacles in the lower GI tract. This observation indicates that *Chlamydia* may depend on plasmids for its dissemination to the large intestine, while chromosome-encoded factors are necessary for sustaining colonization(18). The plasmid-encoded protein Pgp3 is reported to be essential for Chlamydia’s ability to withstand the acidic conditions encountered in the stomach and to bypass the CD4+ T cell barriers of the small intestine. However, once Chlamydia arrives at the large intestine, the role of Pgp3 diminishes. Instead, chromosome-encoded open reading frames TC0237 and TC0668 become crucial for *Chlamydia* to escape the influence of IFN-γ produced by group 3-like innate lymphoid cells in the large intestine(20, 21). Despite the presence of various pathogenic virulence factors of *Chlamydia*, additional research is warranted to further investigate the interactions between Chlamydia and the host within the gut. This is particularly important as the immune responses and the defensive mechanisms of gut epithelial tissues can be notably affected by gut microbiota and metabolites(22), yet related studies remain limited in the current literature. *C. muridarum* infection of the genital tract in mice is one of the most widely used preclinical models for examining chlamydial pathogenesis and assessing chlamydial vaccines(23–25). Following intravaginal inoculation, *C. muridarum* is known to lead to hydrosalpinx and infertility in mice, closely replicating the tubal adhesions seen in women during laparoscopy(26, 27). Beyond infecting the mucosal surfaces of the mouse genital and respiratory tracts, *C. muridarum* can also inhabit the GI tract for prolonged periods without causing significant pathological changes in the genital region(12).

The ability of *Chlamydia* to efficiently colonize the gut creates new opportunities for developing a gut-based vaccination strategy. Among various vaccination approaches, live *Chlamydia* inoculation has demonstrated effectiveness, but with associated safety concerns(28). One major concern is the potential for live *Chlamydia* to facilitate the transmission of the pathogen. Although the specific roles of *Chlamydial* species within the GI tract remain largely undefined, recent investigations using oral inoculation with *C. muridarum* has been proven to trigger strong transmucosal immunity against subsequent chlamydial infections in both genital and respiratory systems, indicating the feasibility of developing *C. muridarum* as an oral vaccine for stimulating transmucosal immunity. The implementation of oral inoculation with *C. muridarum* as a vaccination strategy extends beyond mere replication of human vaccination process(29). Reports suggest that gut inoculated *C. muridarum* can induce cross-species immunity against human genital Chlamydia infections as mice with previous exposure to *C. muridarum* have demonstrated heterotypic protection against later infections involving *C. trachomatis*, which implies that the mouse-adapted *C. muridarum* could potentially protect humans from *C. trachomatis* infections(30). As a result, research teams are currently focused on developing *C. muridarum* derived vaccines for human application, and our model has the potential to yield valuable insights to support this ongoing effort(31, 32).

Our mouse model indicated that live Chlamydia elicited a stronger protective response compared to inactivated Chlamydia. As shown in Figure 1, live *Chlamydia* could consistently recover from the gut during the immunization. Given the uncertain nature of gut *Chlamydia* infections, researchers are investigating mutant Chlamydia as an attenuated live vaccine option to elicit a protective immune response while minimizing the pathogenic effects associated with gut colonization(31). In this study, we established a time series following gut *Chlamydia* inoculation, analyzing microbial, metabolic, and immune responses post-immunization. Our findings show that gut *Chlamydia* infections induce distinct changes in microbial and metabolite profiles, resulting in increased antigen-specific antibody production. This work contributes valuable information regarding gut *Chlamydia* infections and vaccine development.

Previous studies have suggested that *Chlamydia* colonization in the gut is nonpathogenic, as assessed through macroscopic evaluations and H&E staining(9, 33). However, our transmission electron microscopy observations of gut epithelial ultra-structures revealed significant alterations in gut tissue as late as day 30 post-inoculation in the live *Chlamydia* group (Fig. 2 c1, c2). Specifically, we observed a reduction in the number of microvilli, a shortening of their length, and swelling of mitochondria within the cytoplasm. These findings raise important questions about the safety and potential impacts of wild-type live *Chlamydia* as a vaccine, underscoring the necessity for further research in this area. Our results support the development of mutant strains of *Chlamydia* with reduced virulence for use in future vaccine strategies and underscore the necessity for evaluating ultra-structures.

Notably, while *Chlamydia* could consistently be detected in the gut via rectal swabs in the live *Chlamydia* group (Fig. 1 A), it was only detectable on day 7 through 16S sequencing when rectal shedding levels were elevated (Fig. 6 C). As an intracellular pathogen, *Chlamydia* primarily resides within infected epithelial cells, and fecal samples usually contain fewer shedding cells compared to rectal swabs. This suggests that feces may not be the optimal sample for detecting gut *Chlamydia* infections(11).

The observed alterations in gut microbial diversity and metabolite composition following vaccination underscore the importance of the gut microbiome and metabolites as key mediators of vaccine efficacy(34). The identification of 129 distinct microbial taxa and 186 metabolites illustrates the complex biochemical environment influenced by *Chlamydia* vaccination, suggesting that these changes may affect the host’s immune responses and protective effects against subsequent *Chlamydia* exposures (Fig. 7). Notably, certain genera, such as *Lactobacillus*, showed positive correlations with protective antibody production, indicating their potential as biomarkers for vaccine efficacy. In contrast, genera like *AF12* and metabolites such as L-Dopa and PAF C-16 exhibited negative correlations with antibody levels, suggesting that these factors may lead to less favorable immune outcomes (Fig. 8). These findings resonate with existing literature connecting gut microbiota diversity to immune health, emphasizing the need for a holistic approach to vaccine development that considers gut microbiota interactions(35). It is well-documented that live *Chlamydia* provides greater protection compared to inactivated *Chlamydia* vaccines and the underlying reasons for this enhanced efficacy remain to be fully elucidated(4). This study may offer valuable insights that contribute to a better understanding of this phenomenon (Fig. 9).

**Figure 9:**
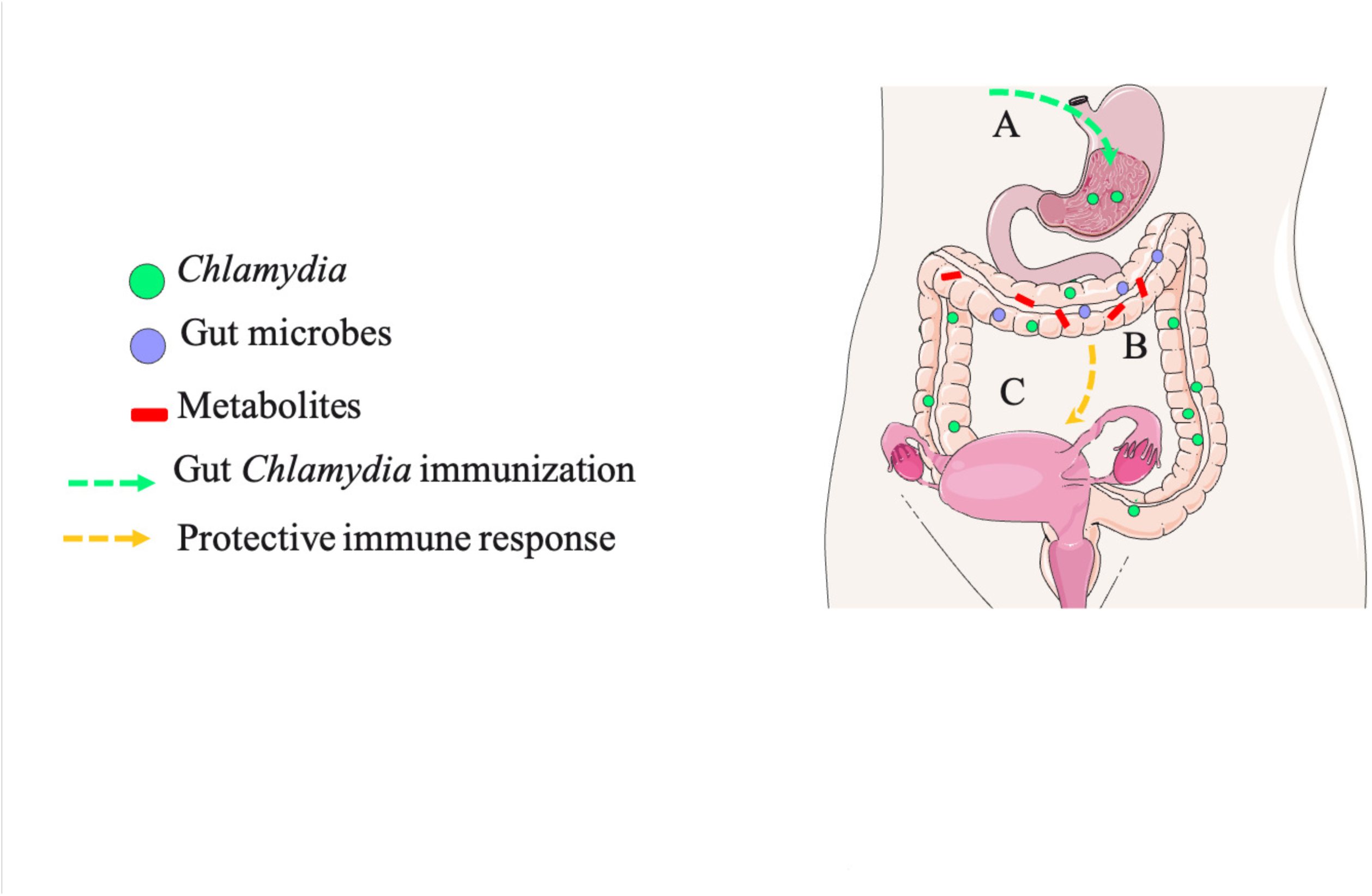
Schematic overview of the impact of oral *Chlamydia* vaccination on the gut microbiome, metabolite composition, and protective immune responses. This hypothetical model illustrates the interactions between oral Chlamydia vaccination, host gut microbiome, and metabolite composition. [A] Inoculation of *Chlamydia* into the GI tract leads to Chlamydial colonization within the GI environment. [B] Gut *Chlamydia* induces changes in microbial and metabolite profiles, which may regulate the host’s protective immune response against Chlamydia. [C] *Chlamydia* specific transmucosal immune effects on the host are depicted. Portions of the figure utilize images from Servier Medical Art, licensed under the Creative Commons Attribution 3.0 Unported License

Despite these insights, this study has limitations that should be acknowledged. Firstly, while we focused on gut microbiome and metabolite analysis, we did not conduct comprehensive serological metabolomics, which limits our understanding of systemic metabolic changes and their potential correlations with the gut environment and immune responses. Inclusion of serum metabolite profiling could provide a more complete picture of how oral vaccination influences host metabolism.

Furthermore, although we identified several potential biomarkers associated with vaccination efficacy, our findings are primarily rooted in observed changes in gut microbiota, metabolite alterations, and subsequent mathematical analyses. Further validation through dedicated animal studies will be essential to confirm these biomarkers and establish their biological relevance. Without animal model confirmation, it remains uncertain whether these correlations will hold true in broader biological contexts.

This study provides information for the interplay between oral *Chlamydia* vaccination, gut microbiome alterations, and gut metabolite profiles. The implications extend beyond *Chlamydia* vaccination alone; the nature of oral live *Chlamydia* vaccination enables prolonged colonization in the gut, yielding important insights into *Chlamydia* persistence in this niche. This research enhances our understanding of the biological significance of gut colonization and improves our grasp of host-*Chlamydia* interactions. By elucidating these dynamics, our study may inform strategies for managing Chlamydia gut infections and the development of more effective approaches for detecting and preventing these infections in the future(3).

## MATERIALS AND METHODS

### Animals

C57BL/6J female mice, aged 5-6 weeks, were sourced from Vital River in Beijing and maintained under controlled environmental conditions (temperature: 22 °C; light/dark cycle: 12 hours each). Following a one-week acclimatization period in the laboratory, these mice were used for experiments that received approval from the Ethics Committee of the Institute of Pasteur at the Shanghai Academy of Sciences, adhering to the guidelines set by the Chinese Council on Animal Care. The mice were randomly arragened into 3 groups, Control groups (labeled as C) and De_*Chlamydia* group (labeled as J) and *Chlamydia* group (labeled as M).

### Preparation of *Chlamydial* Organisms

All *C. muridarum* clones utilized in this study were obtained from the Nigg3 strain (GenBank accession number CP009760.1). The luciferase-expressing clone of *C. muridarum* has been previously documented(36). *Chlamydia* was propagated in HeLa cells and subsequently purified into EBs as outlined in earlier studies(37). These purified EBs were aliquoted and stored at -80 °C until required. To deactivate the EBs, they were heat-treated at 56 °C for 30 minutes.

### Immunization, Challenge Infections, and Assessment of Genital Pathology

To immunize the mice, purified *C. muridarum* EBs were intragastrically administered to female mice aged 6 to 7 weeks, reflecting the procedures described previously(33). Each mouse received an inoculation of 2 × 10^5 inclusion-forming units (IFUs) of either live or deactivated *C. muridarum*, mimicking oral immunization. The control group was administered a sucrose-phosphate-glutamic acid (SPG) vehicle. Following each inoculation, vaginal and rectal swabs were collected periodically to monitor the colonization of viable *C. muridarum* in the gut and genital tract. Additionally, fecal samples from each group were preserved in liquid nitrogen for further experiments, and serum samples were collected at various time points.

Fifty days post-immunization, the mice were challenged intravaginally with 2 × 10^5 IFUs of *C. muridarum*, and shedding from the genital tract and gut was monitored through swabs. The mice were sacrificed on day 56 post-challenge for evaluation of genital pathologies(9). Furthermore, four mice from each group were sacrificed on day 30 post-immunization to examine gut epithelial structures using transmission electron microscopy, with a focus on hydrosalpinx in the upper genital tract. High-resolution digital photography was employed to document the oviduct hydrosalpinx, which was subsequently scored for severity and incidence within each group.

### Monitoring of *Chlamydia* Infections

To monitor the shedding of live organisms, cervicovaginal and anorectal swabs were collected every 3 to 4 days for the first week, followed by weekly collections. Each swab was immersed in 0.5 ml of SPG and vortexed with glass beads to extract *Chlamydial* organisms, which were then titrated on HeLa cell monolayers in duplicate. The infected cultures were processed for immunofluorescence assays as described below. Inclusions were counted in five random fields per coverslip under a fluorescence microscope. For coverslips with fewer than one IFU per field, the entire coverslip was counted. Coverslips exhibiting significant cytotoxicity in HeLa cells were excluded from analysis.

The total number of IFUs per swab was calculated based on the average IFUs per view, the area ratio of the view to that of the well, dilution factors, and inoculation volumes. If applicable, mean IFUs per swab were derived from serially diluted and duplicate samples. The total count of IFUs per swab was converted to log10 and used to compute the mean and standard deviation for the mice within each group at each time point.

For live imaging, mice infected with the luciferase-expressing *C. muridarum* clone (G5-pGFP-Luci) were imaged using the Xenogen IVIS imaging system (PerkinElmer, Hopkinton, MA) on various days after infection. Prior to imaging, 500 µl of D-luciferin (40 mg/ml in sterile phosphate-buffered saline [PBS]) was injected intraperitoneally into each mouse. Twenty-five minutes after injection, the mice were anesthetized with 2% isoflurane. Bioluminescent images of the entire mouse were captured as described previously(38).

### ELISA for Measurement of Mouse Fecal IgA and Serum IgG Antibodies

To assess IgA antibodies, fecal samples from each mouse were resuspended in PBS solution to achieve a final concentration of 1 mg/10 µl. After centrifugation at 13,000 rpm for 5 minutes, the supernatants were applied neat or diluted 2-fold to 96-well plates pre-coated with purified *C. muridarum* EBs. IgA binding was detected using goat anti-mouse IgA conjugated with horseradish peroxidase (HRP; catalog number 626720; Invitrogen, Waltham, MA) along with a soluble substrate, ABTS (2,2’-azinobis [3-ethylbenzothiazoline-6-sulfonic acid] diammonium salt; catalog number 30931670; Sigma-Aldrich, St. Louis, MO). Absorbance readings were taken at 405 nm using a Synergy H4 microplate reader (BioTek, Winooski, VT), with results expressed as raw optical density (OD) values. For IgG detection, serum samples collected from the tail vein of mice were subjected to a 4-fold serial dilution starting at 1:1,600. IgG binding to plate-coated EBs was identified using a goat anti-mouse IgG-HRP conjugate (catalog number 31430; Thermo Fisher Scientific) as previously described. The same ELISA protocol was applied for IgG isotyping, with serum samples diluted 1:1,600 on *C. muridarum*-coated plates. Results were also reported as raw OD values.

### DNA Extraction and 16S rRNA Gene Amplicon Sequencing

Genomic DNA extraction was performed using the OMEGA Soil DNA Kit (M5635-02, Omega Bio-Tek, Norcross, GA, USA), in accordance with the manufacturer’s instructions, and samples were stored at -20 °C prior to analysis. The quality and quantity of the extracted DNA were evaluated using a NanoDrop NC2000 spectrophotometer (Thermo Fisher Scientific, Waltham, MA, USA) and agarose gel electrophoresis, respectively.

PCR amplification of the V3–V4 region of bacterial 16S rRNA genes was conducted using the forward primer 338F (5’-ACTCCTACGGGAGGCAGCA-3’) and reverse primer 806R (5’-GGACTACHVGGGTWTCTAAT-3’). Sample-specific 7-bp barcodes were included in the primers for multiplex sequencing. Each PCR reaction contained 5 μl of 5× buffer, 0.25 μl of Fast Pfu DNA Polymerase (5U/μl), 2 μl of 2.5 mM dNTPs, 1 μl (10 µM) of each primer, 1 μl of DNA template, and 14.75 μl of ddH2O. The thermal cycling protocol included an initial denaturation at 98 °C for 5 minutes, followed by 25 cycles of denaturation (98 °C for 30 s), annealing (53 °C for 30 s), and extension (72 °C for 45 s), capped with a final extension at 72 °C for 5 minutes. PCR amplicons were purified using Vazyme VAHTSTM DNA Clean Beads (Vazyme, Nanjing, China) and quantified with the Quant-iT PicoGreen dsDNA Assay Kit (Invitrogen, Carlsbad, CA, USA). Following quantification, amplicons were pooled in equal concentrations, and pair-end 2×250 bp sequencing was executed on the Illumina NovaSeq platform with the NovaSeq 6000 SP Reagent Kit (500 cycles) at Suzhou PANOMIX Biomedical Tech Co., Ltd. Sequence data analyses were primarily completed using QIIME2 and R packages (v3.2.0). Alpha diversity indices at the ASV level, such as Chao1 richness estimator, observed species, Shannon diversity index, and Simpson index, were calculated from the ASV table in QIIME2 and visualized as box plots (Fig. S1). ASV-level ranked abundance curves were generated to assess richness and evenness across samples. Beta diversity analyses examined the structural variation of microbial communities among samples, the correlation among samples were analyzed (Fig. S2), and using Bray-Curtis and UniFrac distance metrics, with results visualized through principal coordinate analysis (PCoA). Principal component analysis (PCA) was also performed on genus-level compositional profiles. Taxonomic compositions and abundances were visualized, and Linear discriminant analysis effect size (LEfSe) was applied to identify differentially abundant taxa across groups. Clustering tree analysis of various samples were attached (Fig. S3)

### LC-MS Analysis

LC analysis was performed using a Vanquish UHPLC System (Thermo Fisher Scientific, USA). Chromatography was conducted with an ACQUITY UPLC® HSS T3 column (2.1×100 mm, 1.8 µm; Waters, Milford, MA, USA) in both positive and negative ion modes(39). Mass spectrometric detection of metabolites utilized an Orbitrap Exploris 120 (Thermo Fisher Scientific, USA) with an ESI ion source, employing simultaneous MS1 and MS/MS acquisition in data-dependent MS/MS (Full MS-ddMS2) mode. The instrument parameters included a sheath gas pressure of 40 arb, an auxiliary gas flow of 10 arb, and spray voltages of 3.50 kV (ESI+) and -2.50 kV (ESI-). Capillary temperature was maintained at 325 °C, with a mass range of m/z 100-1000 and an MS1 resolving power of 60,000 FWHM. The number of data-dependent scans per cycle was set to 4, with an MS/MS resolving power of 15,000(40). The Mfuzz method was employed for cluster analysis of metabolite patterns across different samples, utilizing a fuzzy c-means algorithm to establish relationships between time and metabolites across clusters(41).

### Statistical Analysis

Quantitative data, including viable organism counts (IFUs) and genome copies, were analyzed using the area under the curve (AUC) method, the Wilcoxon rank-sum test and Kruskal-Wallis test. For qualitative data, such as incidence rates, Fisher’s exact test was employed. Additionally, semiquantitative data were also analyzed using the Wilcoxon rank-sum test. A significance threshold of *p* < 0.05 was set to determine statistical significance.

## Conflict of Interest

The authors declare that the research was conducted in the absence of any commercial or financial relationships that could be construed as a potential conflict of interest.

## Author Contributions

YYH, JW and CQS wrote the original draft. JW, QZ, ZZZ, and XS reviewed and edited the manuscript. YYH and TYZ contributed to the figure preparations. QT, TYZ and LYW provided the conceptualization and funding. All authors contributed to the article and approved the submitted version.

## Funding

This research is supported by National Natural Science Foundation of China for the Youth (32100162) to QT and (32000138) to TZ. The Natural Science Foundation of Hunan Province, China (2022JJ70092 and 2023JJ40355), the Science and Technology Innovation Program of Hunan Province (2024RC3233 and 2021SK4021).

## Acknowledgments

We express our gratitude to Prof. Wu Xiang from the Department of Parasitology at Central South University’s Xiangya Medical School for her support in the preparation of *Chlamydia*. Additionally, we extend our thanks to Suzhou PANOMIX Biomedical Tech Co., Ltd for their expertise in conducting the 16S rRNA sequencing and LC-MS analysis, which were crucial for our research.

## Data Availability Statement

The original contributions presented in the study are included in the article/supplementary material, further inquiries can be directed to the corresponding authors.

